# Epigenetic clocks and inflammaging: pitfalls caused by ignoring cell-type heterogeneity

**DOI:** 10.1101/2025.01.21.634004

**Authors:** Xiaolong Guo, Andrew E. Teschendorff

**Affiliations:** CAS Key Laboratory of Computational Biology, CAS-MPG Partner Institute for Computational Biology, 320 Yue Yang Road, Shanghai 200031, China

## Abstract

Epigenetic clocks are machine learning predictors of chronological age that have consistently been shown to be also informative of biological age measures, such as all-cause mortality. A recent study has argued against the use of machine learning methods for building epigenetic clocks, on grounds that these predictors do not enrich for features that correlate with disease. In particular, it argues that standard epigenetic clocks can’t measure inflammaging, an age-related trait, proposing an ad-hoc ‘feature rectification’ strategy and an ‘inflammation clock’ that predicts age-acceleration in inflammatory diseases. Here we demonstrate that this inflammation clock only captures an increase in the neutrophil to lymphocyte ratio associated with inflammatory diseases like rheumatoid arthritis, and that it fails to capture the shift from naïve to mature lymphocytes, which is a key feature of inflammaging. As such, their clock does not measure inflammaging, and is subsumed and outperformed by an ordinary cell-type deconvolution algorithm. Our analysis underscores the critical importance for epigenetic clock and epigenome-wide association studies to always estimate cell-type fractions in the tissue of consideration, and to adjust for its variation, in order to disentangle effects due to changing cell-type composition from DNA methylation changes that happen in specific cell-types.

---

The epigenome-wide association study (EWAS) field is now over a decade old ^1^. The aim of these EWASs is to identify differentially methylated cytosines (DMCs) associated with a wide range of exposures and phenotypes, including aging. Overwhelmingly, the tissue of choice has been whole blood (WB) or peripheral blood mononuclear cells (PBMC). These are complex tissues composed of many different immune cell types, each one characterized by a unique DNA methylation (DNAm) profile ^2-5^. Some of the earliest EWASs already noted the importance of taking cell-type heterogeneity (CTH) into account when identifying differentially methylated cytosines (DMCs) ^6,7^. For instance, one of the earliest EWAS by Liu et al studied Rheumatoid Arthritis (RA), an inflammatory auto-immune disease, demonstrating how the overwhelming majority of RA-associated DMCs merely reflect the increase in the neutrophil to lymphocyte ratio that is associated with a condition like RA ^6^ (**Fig.1a-b**). Indeed, most of these DMCs disappear upon adjustment for this shift in the neutrophil to lymphocyte ratio (**Fig.1b**). In other words, most of the original RA-DMCs do not reflect DNAm changes that happen in specific immune cell-types of RA cases compared to controls. Thus, it is perplexing how a recent study by Skinner and Conboy ^8^ does not adjust for this underlying CTH when analyzing the exact same EWAS by Liu et al; this despite numerous articles that have over the last decade continued to emphasize the critical need to estimate immune cell-type fractions in blood and to adjust for this variation ^9-12^. Recent articles have also highlighted the importance of considering CTH when building and interpreting epigenetic clocks ^13-16^. Hence, it is specially perplexing how Skinner and Conboy don’t acknowledge cell-type heterogeneity even once in their whole manuscript, i.e. no biological or statistical justification is given for ignoring CTH.

**Fig 1:**
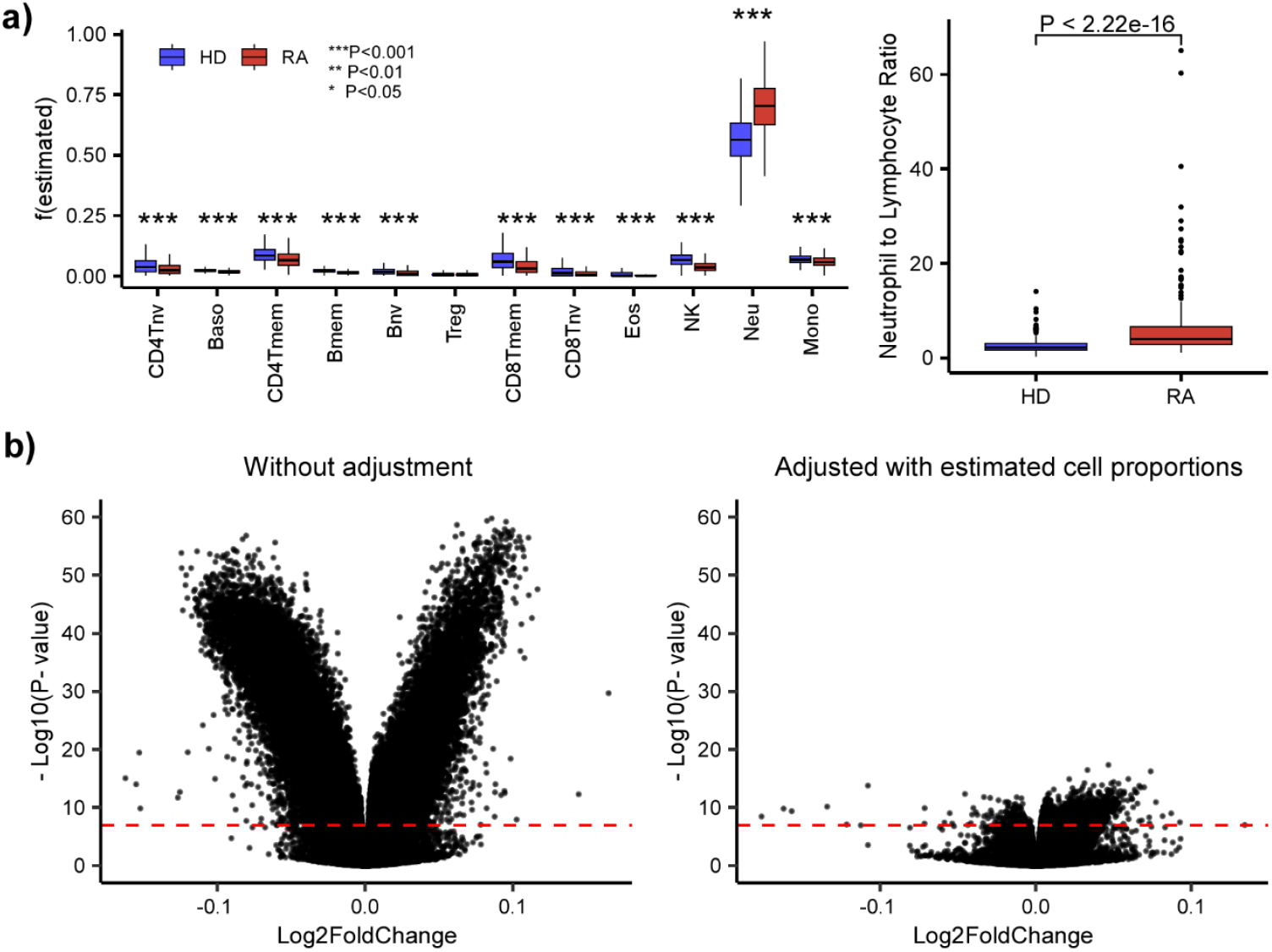
Rheumatoid Arthritis (RA) is associated with many DMCs that reflect a shift in the neutrophil to lymphocyte ratio. **a)** Boxplots of estimated immune cell fractions in the Liu et al DNAm dataset of RA cases and healthy controls (HD), with the fractions as estimated using EpiDISH and a DNAm reference panel for 12 immune cell-types. P-values computed using a two-sided Wilcoxon rank sum test. Right boxplot compares the neutrophil to lymphocyte ratio. **b) Left:** Volcano plot for RA Liu et al EWAS without adjustment for cell-type heterogeneity. CpGs above the red dashed line (Bonferroni threshold) constitute RA-DMCs. **Right:** Same as left but after adjustment for the 12 immune cell-type fractions. Thus, the RA-DMCs that pass the threshold correlate with RA independently of changes in immune cell-type composition.

In their work ^8^, Skinner and Conboy question the value of current epigenetic clocks that have been derived using penalized multivariate regression (Elastic Net) ^17-19^, on the basis that they don’t predict age-acceleration in an inflammatory condition like RA. Instead, they propose a ‘feature rectification’ strategy to derive an ‘inflammation-clock’ (InflClock) that they claim measures inflammaging ^20-23^. However, as described above, their clock was derived without adjustment for CTH. Consequently, all their clock is likely to measure is the shift in the neutrophil to lymphocyte fractions, which is not necessarily specific to inflammaging per se ^20-23^. To demonstrate this, we applied their InflClock to 15 whole blood datasets, whilst also estimating fractions for 12 immune cell-types (neutrophils, monocytes, eosinophils, basophils, natural killer cells, memory and naïve B-cells, memory and naïve CD4+ T-cells, memory and naïve CD8+ T-cells and T-regulatory cells) using our EpiDISH framework ^24,25^. We found a very strong association of the InflClock with increased neutrophil and decreased lymphocyte fractions in each of the 15 cohorts (**Fig.2**).

**Fig 2:**
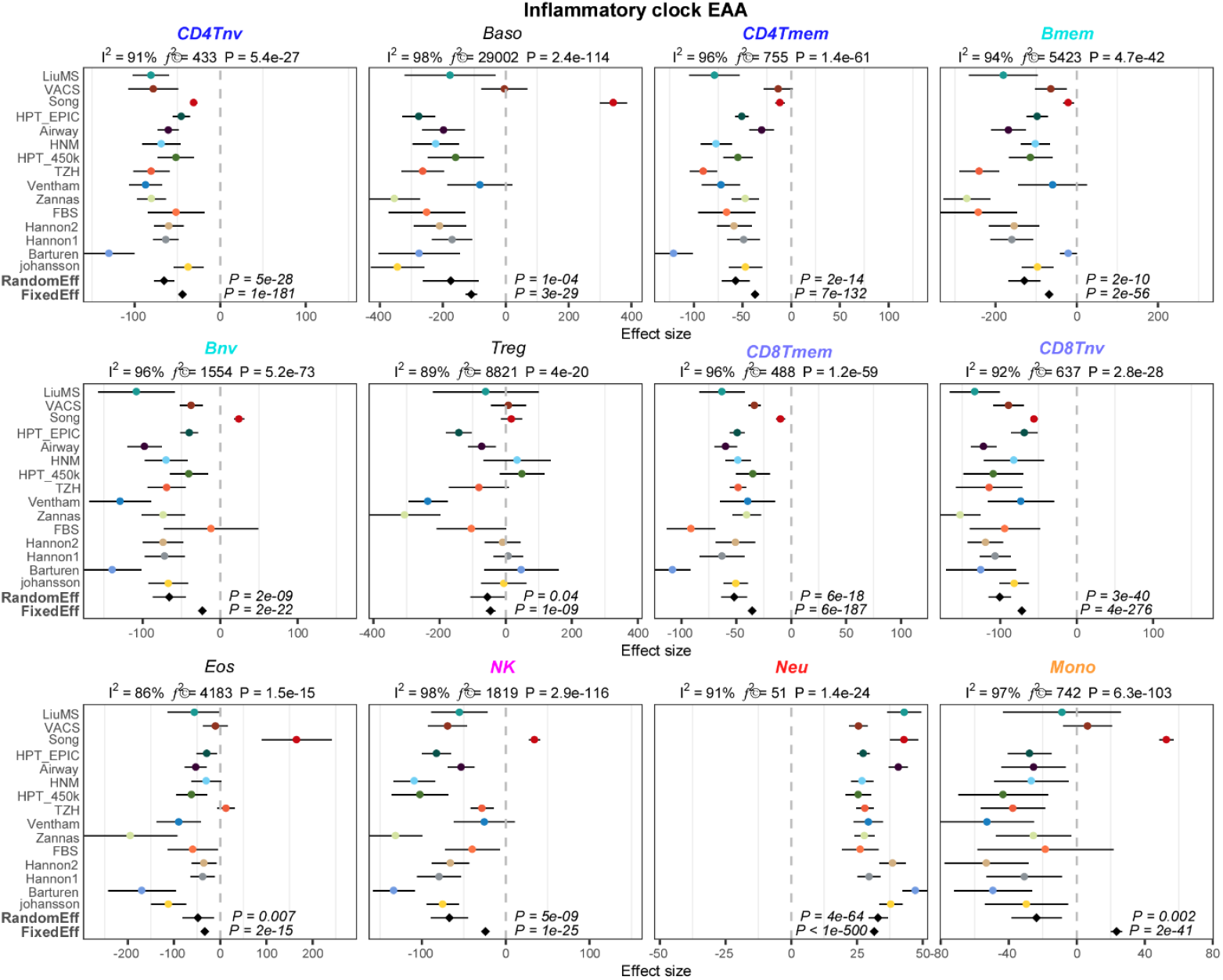
Forest plots of association between the InflClock’s EAA and 12 immune cell-type fractions. Each panel is a forest plot between an immune cell-type’s estimated fraction and its association with the InflClock’s extrinsic age acceleration (EAA), as assessed over 15 independent whole blood cohorts. For each Forest plot, we provide the meta-analysis P-value for a random effect and fixed effect models. Error bars represent 95% confidence intervals for the estimates. On the top of each panel we also provide the heterogeneity indices and associated P-values. Immune cell-types are abbreviated as: CD4Tnv=naïve CD4+ T-cells, CD4Tmem=memory CD4+ T-cells, Bnv=naïve B-cells, Bmem=memory B-cells, CD8Tnv=naïve CD8+ T-cells, CD8Tmem=memory CD8+ T-cells, Baso=basophils, Eos=eosinophils, Treg=T regulatory cells, Mono=monocytes, NK=natural killer cells, Neu=neutrophils.

Hence, this questions the validity of InflClock, as it merely recapitulates the variation in the neutrophil to lymphocyte ratio, which could vary due to a whole plethora of factors (including notably acute inflammation), not just chronic low-grade inflammation. Indeed, among the 15 whole blood datasets analyzed, there are many that involve cohorts of reasonably healthy individuals, for instance the Airway ^26^, Hannum (HNM)^27^, Taizhou (TZH) (You et al) ^28^ and Johansson ^29^ cohorts are not enriched for individuals suffering from an inflammatory disease like RA or IBD. Importantly, inflammaging would also be associated with shifts within the lymphocyte compartment, in particular one would expect the InflClock to correlate with an increase in the memory T-cell fractions and to anti-correlate with the corresponding naïve T-cell fractions ^30-33^. Yet the meta-analysis over the 15 cohorts unequivocally demonstrates that both memory and naïve T-cell fractions correlate negatively with the InflClock (**Fig.2**), further supporting the view that InflClock is only capturing an increased global neutrophil-to-lymphocyte ratio.

To further demonstrate this, we built a separate inflammation score from a large meta-analysis of EWASs, where DNAm was correlated to levels of C-reactive protein (CRP), a well-known marker of inflammation including low-grade chronic inflammation ^34^ (**Methods**). Specifically, we used 1765 CpGs whose DNAm levels are robustly correlated to CRP across over 22,000 whole blood samples, to build a sample specific inflammation score predictor (InflScore). When applied to the same 15 whole blood cohorts from our earlier meta-analysis, this confirmed that InflScore captures shifts in the naïve to mature T-cell fractions, specifically an increase in mature T-cell fractions compared to naïve ones, with a corresponding increase in NK-cells, whilst the neutrophil proportions decreased (**Fig.3**). In other words, the InflScore and InflClock are measuring completely different processes, yet Skinner and Conboy claim that InflClock measures inflammaging.

**Fig 3:**
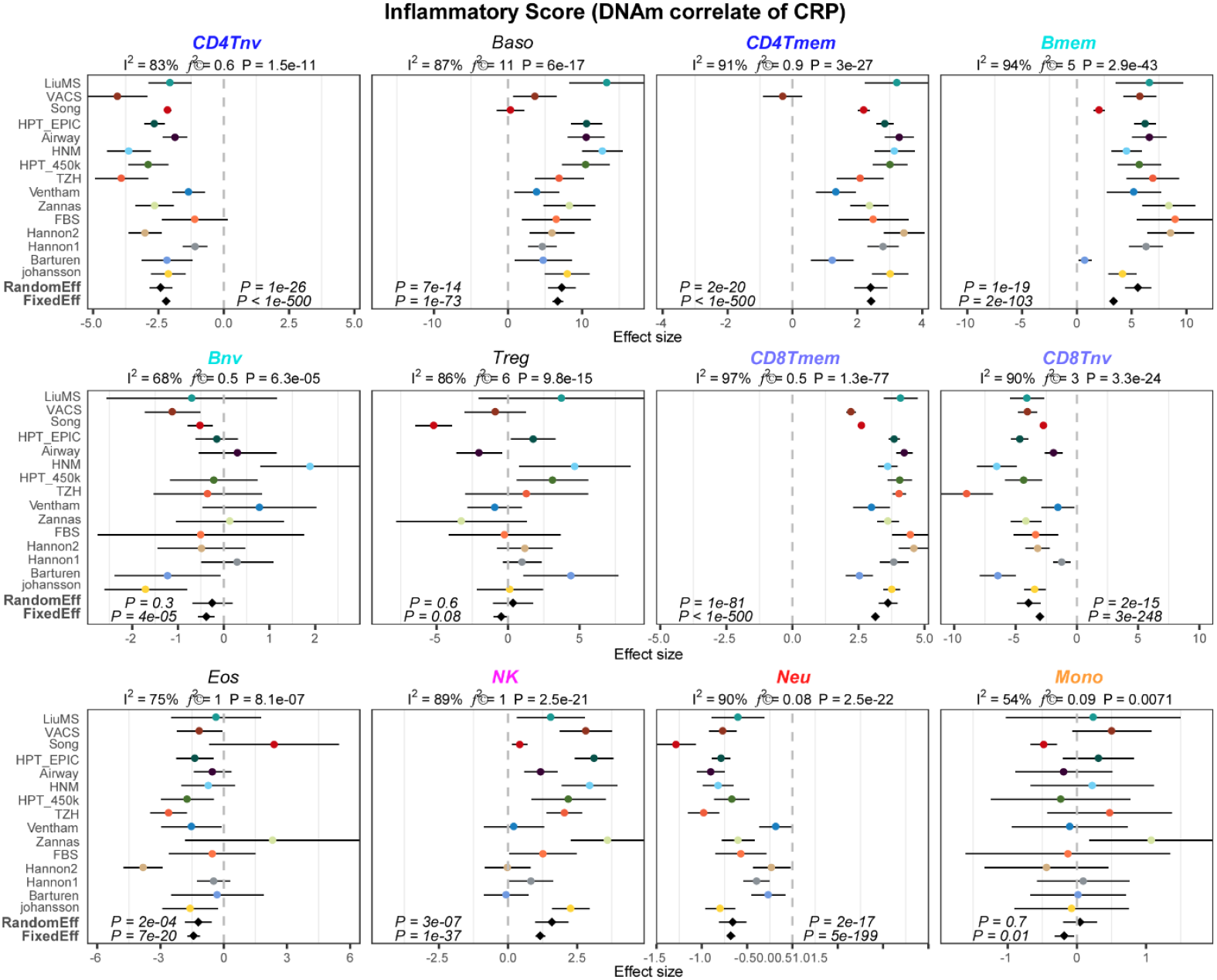
Forest plots of association between the Inflammation Score and 12 immune cell-type fractions. Each panel is a forest plot between an immune cell-type’s estimated fraction and its association with an Inflammation Score which is a DNAm proxy for CRP levels, as assessed over 15 independent whole blood cohorts. For each Forest plot, we provide the meta-analysis P-value for a random effect and fixed effect models. Error bars represent 95% confidence intervals for the estimates. On the top of each panel we also provide the heterogeneity indices and associated P-values. Immune cell-types are abbreviated as: CD4Tnv=naïve CD4+ T-cells, CD4Tmem=memory CD4+ T-cells, Bnv=naïve B-cells, Bmem=memory B-cells, CD8Tnv=naïve CD8+ T-cells, CD8Tmem=memory CD8+ T-cells, Baso=basophils, Eos=eosinophils, Treg=T regulatory cells, Mono=monocytes, NK=natural killer cells, Neu=neutrophils.

Incidentally, the 1765 CpGs associated with CRP levels were derived by adjusting for CTH but only at the coarse resolution of 7 immune cell-types ^34^, which did not include separate naïve and memory T-cell components. This explains why the InflScore is capturing the increased memory and decreased naïve T-cell fractions, once again highlighting the need to adjust for CTH at a higher resolution if the aim is to identify DNAm changes in specific cell-types that are associated with chronic low-grade inflammation. In this regard it should be further noted that Skinner and Conboy do not demonstrate that their InflClock captures DNAm changes within neutrophil, lymphocyte or monocyte subsets that reflect an underlying inflammatory state, further undermining their claim that InflClock measures inflammaging ^20-23^.

The failure to estimate immune cell fractions and adjusting for them when deriving the InflClock also seriously undermines the need for InflClock, since the biological information content of this clock is subsumed by the immune cell fractions that are estimated with a cell-type deconvolution algorithm. In other words, we don’t need InflClock, since the cell-type deconvolution algorithm, by virtue of estimating the underlying immune-cell fractions, can yield excellent correlates of inflammatory disease status. For instance, a cell-type deconvolution algorithm such as EpiDISH clearly predicts an increased neutrophil fraction in RA cases compared to controls without the need for any training or machine-learning (**Fig.1**). Moreover, estimated immune cell fractions also vary in relation to other inflammatory conditions, such as multiple sclerosis, systemic lupus erythematosus or psoriasis (**Fig.4**), again without the need for training, machine learning or an ad-hoc ‘feature rectification’ strategy as proposed by Skinner and Conboy. In addition, the cell-type deconvolution algorithm is able to provide insight, demonstrating that it is the ratio of neutrophils to memory CD8+ T-cells that is a common feature of these inflammatory conditions. Furthermore, if the aim is to capture cell-type specific DNAm changes that are associated with inflammation, and which do not merely reflect the changes in immune cell-type composition that accompany this process, one must estimate immune cell fractions and use these fractions as covariates in the supervised linear models ^6,7,9^. To stress this point, the InflClock predictor is uninformative of which CpGs are changing in the individual cell-types of an inflammatory condition.

**Fig 4:**
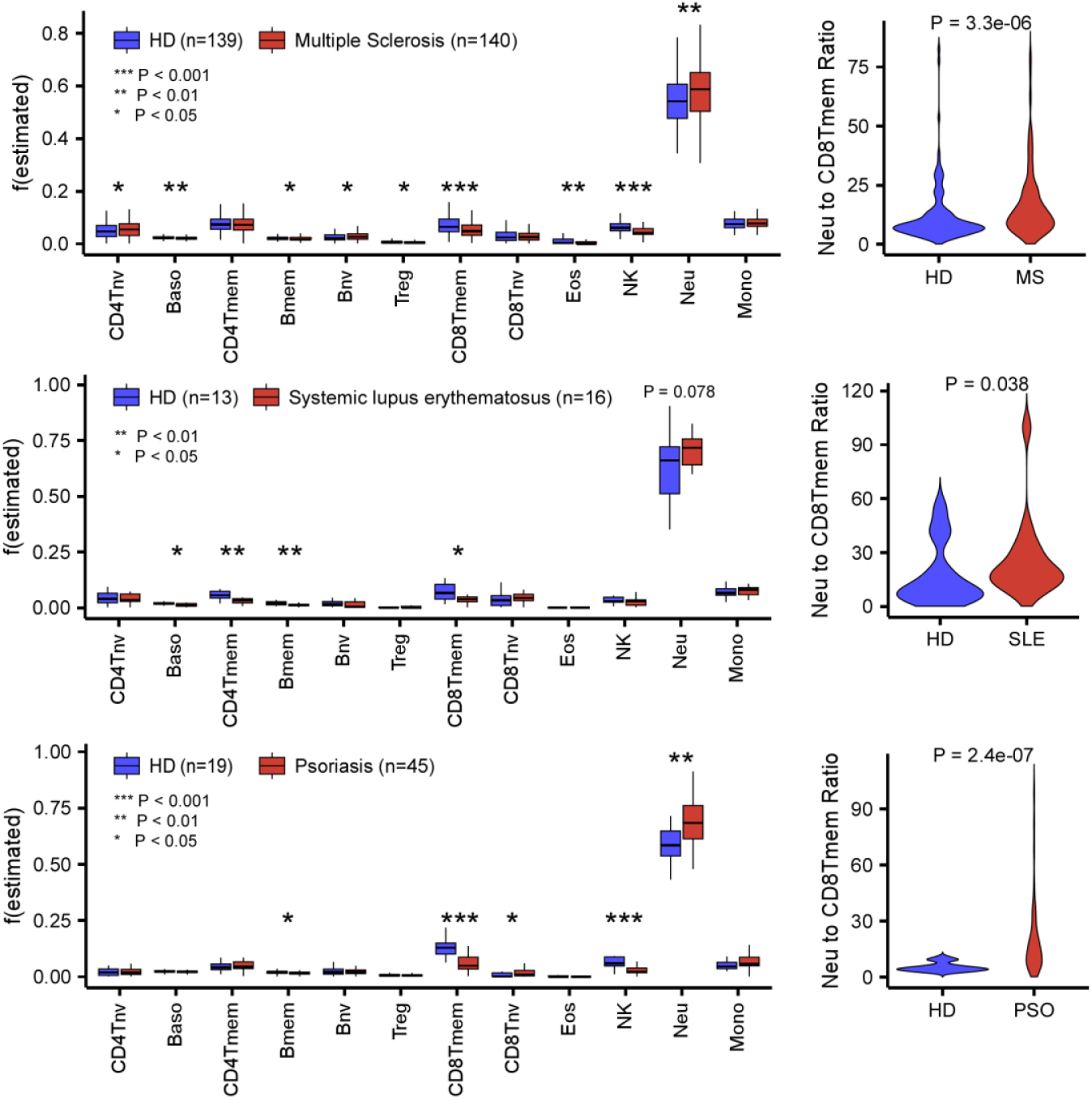
Increased neutrophil to CD8+ T-cell memory ratio characterizes inflammatory conditions. In each of 3 DNAm datasets profiling inflammatory conditions and age-matched healthy donors (HD), boxplots display the corresponding estimated immune cell-fractions for 12 immune cell-types. P-values computed with a two-tailed Wilcoxon rank sum test. Violin plots on the right compare the neutrophil to CD8+ T-cell memory fraction between cases and controls.

The construction of the InflClock itself is also problematic. Skinner and Conboy argue against the use of machine learning methods like penalized regression ^35^ when deriving a clock for say chronological or biological age, on grounds that many of the features in the clock are not themselves strongly associated with age or with age-related biological processes such as inflammation. Rheumatoid Arthritis (RA) is used as an example to illustrate how existing epigenetic clocks fail to predict age-acceleration, when according to Skinner and Conboy this acceleration should be expected. However, we respectfully disagree with their argument, as we now explain.

First of all, the nature of penalized multivariate regression is to remove redundancy among features that strongly correlate with a phenotype of interest, be it age or a disease. Hence, many features that correlate strongly with the phenotype of interest would not necessarily enter the model because their information content is already captured by other CpGs that also correlate strongly with the same phenotype. Moreover, the optimization involved in penalized multivariate regression aims to identify complex linear combinations of CpGs that improve predictive accuracy. As such, specific CpGs may not necessarily be strongly associated with the phenotype as assessed over all training set samples, but may be strongly associated for specific sample subsets. This is more explicit in the context of other machine learning methods like Random Forests ^36^ or even deep-learning ^37^, but still applies, implicitly so, in the case of penalized multivariate regression. Thus, it is perfectly natural for a machine learning method to not include all features strongly associated with the phenotype of interest, and indeed it is generally speaking not recommended to perform Gene Set Enrichment Analyses only on the features that enter such machine learning predictors.

Second, contrary to Skinner and Conboy’s claim, the sign of estimated regression coefficients in a penalized multivariate regression model will in general correlate with the sign of their univariate coefficients. To demonstrate this, we used the Liu et al DNAm dataset to build an Elastic Net predictor for RA. We found that the sign of the coefficients of the selected CpGs in the RA-predictor were strongly associated with the sign of their univariate correlations (**Fig.5a**). To show that this also extends to age as the phenotype, we compared the multivariate regression coefficients of the 353 Horvath clock CpGs to their univariate associations with age in 3 independent cohorts of healthy individuals (the 335 controls from Liu et al ^6^, 710 healthy Han Chinese from You et al ^38^ and 732 healthy individuals from Johansson et al ^29^). In all cases, we observed an excellent correlation between the multivariate and univariate coefficients (**Fig.5b-d**). Thus, penalized multivariate regression is a perfectly valid and powerful method to build predictors for any phenotype ^35^, and has by far been the most popular and successful method in the epigenetic clock field ^16^. In our opinion, there are at least three reasons why Skinner and Conboy reach a different conclusion. First, when comparing multivariate to univariate coefficients they take the absolute value of the clock CpG regression coefficients, comparing them to the R^2^ values derived from univariate regression, yet this discards important information about the directionality of change and would clearly lead to biased, over pessimistic results. Second, the multivariate clock coefficients (say from Horvath ^17^ or Zhang clocks ^39^) were derived from fairly large training DNAm datasets (typically around 7,000 to 10,000 samples or more), and these are then being compared to univariate coefficients derived from a much smaller composite DNAm dataset encompassing around 1000 samples ^8^. Consequently, the relatively low univariate coefficient values of Horvath clock CpGs in the composite dataset of Skinner and Conboy could be driven in part by low power. Aging specially is associated with small effect sizes, and hence it is well known that very large sample sizes are required to ensure sufficient power, and most importantly, to ensure the stability of feature rankings, a key point that was already made almost 20 years ago in the context of cancer prognosis ^40^. However, as shown in Fig.5, when we compared the elastic net Horvath clock CpG coefficients to their univariate coefficients in 3 DNAm datasets, each encompassing fewer than a 1000 samples, we still observed a remarkably good agreement. Thus, although low power may certainly explain why the correlation is not much stronger, this does not seem to explain Skinner and Conboy’s result. A third and more important reason is that Skinner and Conboy compute univariate correlations with age across a composite DNAm dataset made up of 4 different cohorts, not adjusting for cohort of origin. Given that aging is associated with a small effect size, any batch effect associated with cohort of origin, could have a dramatic confounding effect on these univariate coefficients. Indeed, one should bear in mind that computing simple univariate coefficients is more likely to fit to random technical variation: reason why Horvath’s clock, which was trained on over 7,800 samples from over 50 different studies, is robust to batch effects, is precisely because it was trained with a penalized multivariate regression and cross-validation procedure that avoids fitting to random variation ^41^, including technical variation. In other words, Skinner and Conboy incorrectly attribute the low concordance of multivariate and univariate coefficients to a ‘problem of penalized regression’ when it is precisely the univariate estimates which are unreliable because they were not derived from a cross-validation procedure.

**Fig 5:**
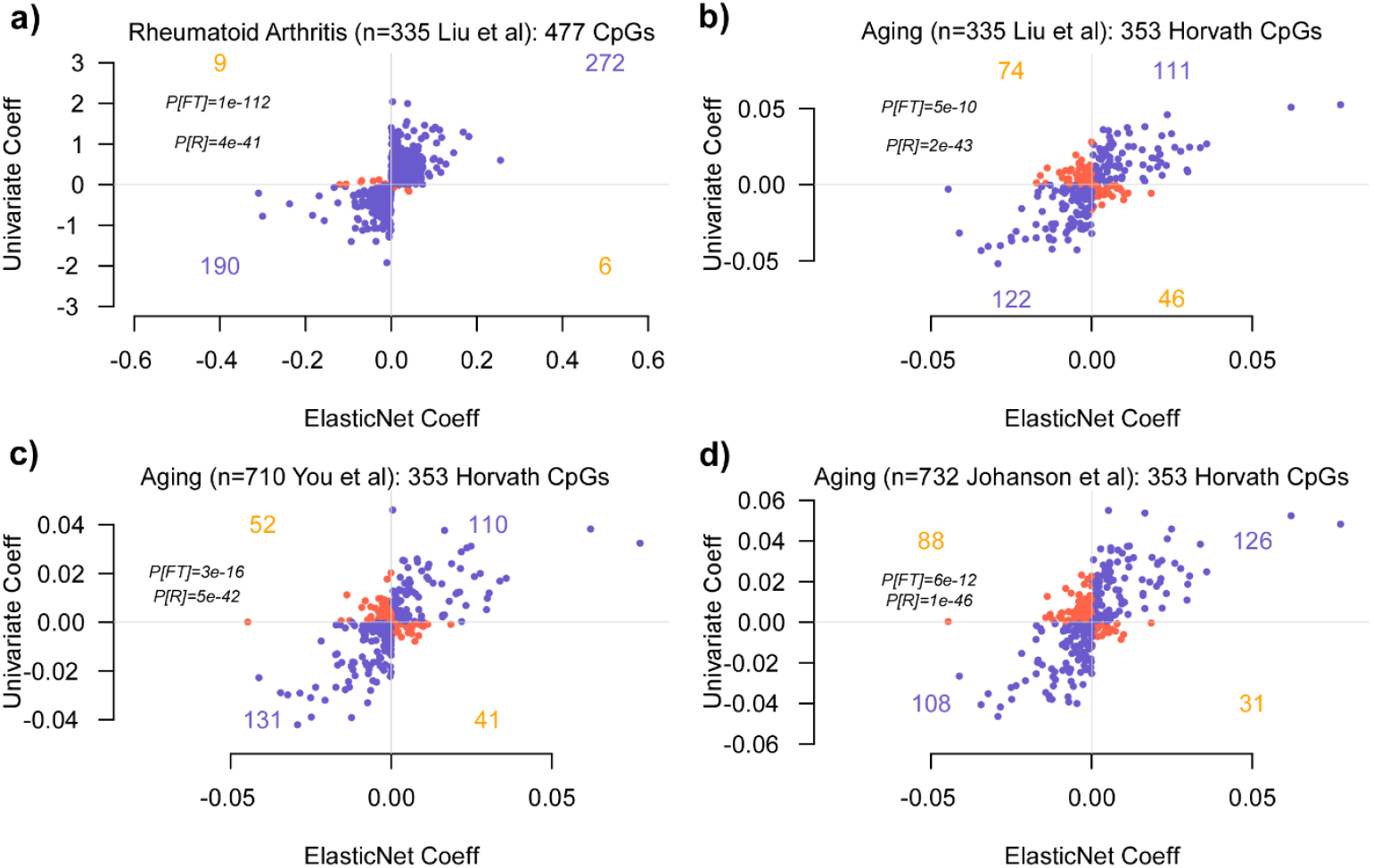
Consistency of multivariate and univariate regression coefficients in association with Rheumatoid Arthritis and aging. **a)** Scatterplot displays the regression coefficient of 477 CpGs with non-zero coefficients in an Elastic Net prediction model for Rheumatoid Arthritis (x-axis) against the corresponding univariate regression coefficients (y-axis). Those CpGs displaying the same directionality are shown in blue, those with inconsistent directionality in red. P-values are derived from a one-tailed Fisher-test (FT) or from a Pearson Correlation test (R). **b)** Scatterplot displays the elastic net regression coefficient of the 353 Horvath Clock CpGs (x-axis) against the corresponding univariate regression coefficients for age, adjusting for sex, smoking and 12 immune cell-type fractions, as evaluated over 335 control samples from the Liu et al cohort (y-axis). Those CpGs displaying the same directionality are shown in blue, those with inconsistent directionality in red. P-values are derived from a one-tailed Fisher-test (FT) or from a Pearson Correlation test (R). **c-d)** As b) but with the univariate CpG coefficients for age computed in the You et al and Johansson et al cohorts. In both cohorts, the regression was adjusted for sex, smoking and 12 immune cell-type fractions.

Third, the expectation of Skinner and Conboy that somehow the signs of coefficients of CpGs in a multivariate predictor for chronological age should be the same as those in relation to a specific condition such as RA is unreasonable and not scientifically sound, as these are inherently different biological processes. Moreover, if the aim of Skinner and Conboy is to build a predictor of RA or inflammation, then there is absolutely no need to consider a clock framework. The biological processes that underpin any given epigenetic clock may or may not be relevant to a particular age-related disease, and it is not scientifically sound to ‘force’ clocks to predict age-acceleration in conditions where they are not meant to be predictive, particularly if the ‘enforcement procedure’ involves a series of ad-hoc steps (i.e. feature rectification) ^8^ that are not based on sound statistical principles. As a concrete analogy, Horvath’s clock is a multi-tissue clock that predicts chronological age across many different tissue-types that differ substantially in terms of their turnover rates (e.g. neurons and colon). As such, Horvath’s clock could never be a mitotic clock, as explicitly demonstrated by previous studies ^42-44^. Following Skinner and Conboy’s argument however, one would naively expect Horvath’s clock to display age acceleration in every cancer-type where incidence increases with age, yet multiple studies have shown that it is not ^42,45^. Unlike mitotic clocks that do display universal age-acceleration across all cancer-types (as expected because of the universal higher proliferation rate of tumor cells)^42-44^, Horvath’s clock does not need to display age-acceleration in cancer since it is not a mitotic clock. Thus, it would not be scientifically sound to ‘alter’ or ‘rectify’ CpG DNAm values of Horvath clock CpGs in order for it to then display universal acceleration in cancer, yet this is exactly what Skinner and Conboy are proposing in the context of inflammation. It is important to understand that there are many different types of epigenetic clocks and that each of these are driven by potentially very different biological processes ^16^, some of which may or may not be relevant to specific diseases or conditions. Thus, the failure of popular epigenetic clocks to predict age-acceleration in a condition such as RA has more to do with these clocks failing to capture the effect of acute inflammation (increased neutrophil to lymphocyte ratio), as opposed to reflecting an inadequacy of a penalized multivariate regression framework. Related to this point, we note that Skinner and Conboy’s claim that epigenetic clocks fail to capture inflammaging is also contradicted by a number of studies reporting such associations ^31,46,47^. As before, the resolution of this conflict hinges on the observation that Skinner and Conboy’s inflammation clock merely captures a signature of acute inflammation (increased neutrophil to lymphocyte ratio) and not ‘inflammaging’ itself.

## Conclusions

As unequivocally demonstrated here, Skinner and Conboy’s inflammation clock and an inflammaging DNAm-based predictor derived from CRP levels, measure completely different biological processes. Hence, the InflClock from Skinner and Conboy does not measure inflammaging. It only captures the known increase in the neutrophil to lymphocyte ratio and is therefore wholly subsumed and outperformed by an ordinary cell-type deconvolution algorithm that can estimate these immune cell fractions without the need for further training or machine learning. Our findings highlight once again how ignoring cell-type heterogeneity in the context of epigenetic clock studies can lead to seriously wrong interpretations and conclusions.

Traditional machine learning methods, in conjunction with cell-type deconvolution algorithms, constitute a powerful combination to build molecular predictors of phenotypes, including chronological age ^39,48^, biological age ^18,49,50^ or inflammatory diseases.

## Methods

### Human adult whole blood DNA methylation datasets

All the cohorts we used were previously normalized and analyzed by Luo et al^24^, including: (1) *LiuMS (GSE106648)*:, 279 HM450k peripheral blood (PB) samples (140 multiple sclerosis patients + 139 controls); (2) *Song (GSE169156)*, 2052 EPIC PB samples; (3) *HPT-EPIC & HPT-450k (GSE210255 & GSE210254)*: 1394 PB samples (EPIC set) and 418 PB samples (450k set); (4) *Barturen (GSE179325)*: 574 EPIC PB samples; (5) *Airwave (GSE147740)*: 1129 EPIC PB samples; (6) *VACS (GSE117860)*: 529 HM450k PB samples; (7) *Ventham (GSE87648)*: 382 HM450k PB samples; (8) *Hannon-1 and 2 (GSE80417 & GSE84727)* : 675 HM450k PB samples and 847 HM450k PB samples for phase-1 and phase-2, respectively; (9) *Zannas (GSE72680)*: 422 HM450k PB samples; (10) *Flanagan/FBS (GSE61151)*: 184 HM450k PB samples; (11) *Johansson (GSE87571)*: 664 HM450k PB samples; (12) *TZH (OEP000260)*: 705 EPIC PB samples; (13) *HNM (GSE40279)*: 656 HM450k PB samples.

### Blood Inflammation Clock (InflClock)

The inflammation clock derives from Skinner and Conboy ^8^, and includes 47 clock CpGs with coefficients all greater than 0. The corresponding coefficients and intercept are provided in the supplementary materials of their paper. The authors trained the clock using age DMCs associated with inflammation, and they adjusted it so that the direction of methylation fraction shifts caused by inflammation aligns with the sign of the model coefficients. Specifically, for an hypermethylated age DMC, if inflammation causes its methylation β value to decrease, it is converted to 1−β. This ensures that the clock, with all positive coefficients, does not have inflammation effects cancel out those caused by age. When applying this clock, it is necessary to use the inflammation signal direction information for these 47 clock CpGs, provided in the supplementary materials, to adjust the dataset being predicted. After predicting the DNAm age, extrinsic age acceleration (EAA) is obtained via the residuals of a linear regression fit of DNAm age versus chronological age.

### Chronic low-grade inflammation score (InflScore)

Wielsher et al. identified 1765 marker CpGs significantly associated with serum C-Reactive protein levels at a Bonferroni threshold (P < 1e−7) in their multi-ethnic meta-analysis ^34^. To apply this signature to other DNAm datasets, we first z-score normalized each CpG across samples in each cohort so that the mean is 0 and the standard deviation is 1. The sign of each CpG’s effect size defines a vector which is then correlated to the z-score normalized DNAm profiles of each sample to yield a sample specific inflammation score (InflScore).

### Estimation of cell-type fractions

In all cohorts, we estimated immune cell fractions using the 12 cell-type DNAm reference matrix for either Illumina EPIC or 450k ^24^, as provided by the EpiDISH Bioconductor R-package. Of note, these DNAm reference panels were built using the DNAm data from Salas et al ^4^ and have been extensively validated by us ^24^. Fractions were estimated using the *epidish* function with the “RPC” method and maxit = 500 ^25^.

### Meta-analyses

In each cohort, associations between the inflammatory clock EAA or inflammation score with immune cell type fractions were assessed using multivariate linear regression. For specific cohorts where additional covariates were available, multivariate regression models with these additional covariates were performed. When regressing EAA to cell type fractions, we always adjusted for sex. When regressing the inflammation score to cell type fractions, we always adjusted for age and sex. Other detailed covariate information should be used for each cohort as described in detail in Luo et al ^24^. For each regression, we extracted the corresponding effect size, standard error, Student’s *t* test, and *P*-value. Then we performed a fixed and random effect inverse variance meta-analysis using the metagen function implemented in the meta R-package (version 7.0-0)^51^.

## Declarations

### Author Contributions

Manuscript was conceived and written by AET. GX and AET performed statistical analyses.

### Ethics approval and consent to participate

Not applicable, as this study only analyses existing publicly available data.

